# Shared predation: positive effects of predator distraction

**DOI:** 10.1101/063230

**Authors:** Mickaël Teixeira Alves, Frédéric Grognard, Vincent Calcagno, Ludovic Mailleret

## Abstract

Simple rules based on population equilibria can characterize indirect interactions in three-species systems but fail to predict them when considering behavioral mechanisms. In this paper, we revisit the effects of shared predation, *i.e*. the situation in which two prey are consumed by a common predator. Such predation usually induces negative indirect interactions between prey, or apparent competition, through an increase of predator density and thus of predation pressure. Two mechanisms can however weaken apparent competition and lead to equivocal signs of indirect interactions. On the one hand, predator distraction, which stems from the difficulty to efficiently forage for different prey at the same moment in time and diminishes the number of prey captured *per* predator. On the other hand, predator negative density dependence limits predator growth. To get further insights into simple rules describing indirect interactions brought about by shared predation, we studied two classes of one-predator-two-prey models exhibiting these two mechanisms. We found robust simple rules derived from predator equilibria which state that at least one prey is favored by the presence of the other when the predators partition their foraging effort between them. These rules thus characterize a surprising wide range of indirect effects including apparent predation, apparent commensalism and apparent mutualism. They also highlight different situations in which larger predator populations do not entail smaller prey populations and in which neither prey species can be negatively affected by the other.

## Introduction

In natural ecosystems, a species interacts with networks of many others. Such a diversity implies a complex set of interactions, which may have major effects on the dynamics of species. This includes higher order effects between species that do not directly interact together, *i.e*. indirect interactions [1]. One of the ecological approaches explored to understand such indirect effects in complex food webs is based on community modules [2]. It consists in the extension of pairwise interactions, as predator-prey interactions, to three or more species systems. In some of these modules, simple rules based on comparison of equilibria can predict the outcome of indirect effects between species, such as the *R*\* rule in one-resource-multi-consumer systems [3] and the *P*\* rule in one-predator-multi-prey systems [4]. These conceptual rules however fail to identify indirect effects in modules that include more complex dynamics induced by behavioral mechanisms.

The introduction of an alternative prey into a one-predator-one-prey system is known to alter the initial interactions through indirect effects [5]. A fairly general result is that this introduction can increase the density of the predator in the long term, which consequently raises the predation pressure on both prey. A well-known example is the decrease of the leafhopper *Erythroneura elegantula*, subsequent to the invasion of the related leafhopper *E. variabilis* in California, which was mediated by a parasitoid common to both species, *Anagrus epos* [6]. Such a long-term negative indirect effect occurring between prey because of the numerical response of a shared enemy has been termed apparent competition [7]. The *P*\* rule states that in this situation the prey that withstands the highest predator equilibrium can lead to the extinction of the other prey through apparent competition [4]. However, the *P*\* rule overlooks the fact that even though predators can forage for many different prey, they can hardly search or forage for all of them simultaneously. Cognitive processes, such as the formation of prey search images [8], or simply the spatial segregation of different prey species, can indeed impose a partition of the time a predator can dedicate to each of its prey. As a predator partitions its foraging time between its prey, it releases its predation pressure on each of them. Such a predator distraction effect, also known as switching, is a widespread mechanism in nature: it has been reviewed in [9] and more recently reported for generalist insect predators [10,11] or parasitoid [12,13], lizards [14] or large mammal predators such as wolves [15] or lions [16]. In the short term, predator distraction tends to induce positive indirect interactions between prey [17]. Positive indirect interactions, termed apparent mutualism in an analogy to apparent competition, can thus undermine predictions based on the *P*\* rule [18,19].

Short-term positive indirect interactions may however be weakened or counterbalanced by the long-term numerical response of the predator, as reported *e.g*. between the thrips *Frankliniella occidentalis* and the whitefly *Trialeurodes vaporariorum* sharing a common predatory mite *Ambliseius swirskii* [20]. Following the introduction of thrips, the predatory mite may relax its predation pressure on each prey in the short term. In the long term however, thrips and whiteflies increase the density of predatory mites and thus eventually experience apparent competition. Therefore, predator distraction and predator numerical response are two important mechanisms in one-predator–multi-prey systems. Since these two mechanisms act on prey density in an opposite manner, co-occurrence may eventually lead to various kinds of indirect effects between prey that we aim here to characterize.

In this paper, we have primarily looked for long-term effects induced by the two previous mechanisms. In a variety of models, it has been shown that apparent competition is prevalent in the long term [7,21]. To our knowledge, only few studies have shown that other kind of indirect effects can occur in the long term, such as apparent mutualism, apparent predation or apparent commensalism. As such, predators and prey that cycle can promote positive indirect effects between prey [22]. When populations do not cycle, mechanisms such as predator satiation, distraction effects, or negative density dependence among predators have been identified as factors promoting the occurrence of positive indirect effects between prey [19]. Here, we concentrated on the influence of the way predators partition their time between their prey, *i.e*. their time partitioning strategy. To this end, we followed Holt’s original definition of indirect effects [7] and evaluated positive indirect effects as the change in a prey equilibrium density in the absence or presence of the other prey. This approach is in line with the *R*\* and *P*\* rules, which are also based on a comparison of equilibria. This also helps interpreting the results, since most experimental studies compare single prey-predator situations to multi-prey–predator ones.

Following [23] to ensure the robustness of our findings, we analyzed two classes of one-predator-two prey models and first considered that predators have fixed preferences for each of their prey, which is termed fixed time partitioning strategy or adaptive switching. We then explored models when predators choose their preferences for each prey regarding the respective density of the latter, which is termed adaptive time partitioning strategy. As a result of the predator distraction effect, we have shown that apparent mutualism between prey always occur in the short term regardless of the time partitioning strategy of the predator. We have also found that, independently of the foraging strategy, indirect effects are always positive on at least one of the prey species in the long term, so that apparent competition never occurs in the proposed framework, but apparent predation, apparent commensalism or apparent mutualism do. Lastly, we noted that time partitioning favors long-term commensalism and mutualism, and that this effect is even stronger if predators adaptively forage. We finally showed that simple rules based on predator equilibria characterize each kind of indirect effects found in this study and discussed the implications of such results with special emphasis on biological control programs.

## Modeling

### A one-predator–one-prey interaction with negative density dependence

The present study is built on the extensions of two classes of predator-prey models with negative density dependence of the predator, which will encompass an alternative prey and foraging preferences. Actually, the fundamental idea behind the analysis of these two classes of predator-prey models is to test the influence of model structure on results, and thus ensure the robustness of simple rules characterizing indirect effects.

Predator-prey interactions are commonly represented with the classical Lotka-Volterra model, which has a significant historical importance in earlier literature about indirect effects [7]. We started the study with a modified Lotka-Volterra model and represented prey growth through a prey-dependent logistic equation, and the predator negative density dependance through a quadratic term in the predator differential equation [19,24]. With *N* and *P* corresponding to prey and predator population densities, the model reads

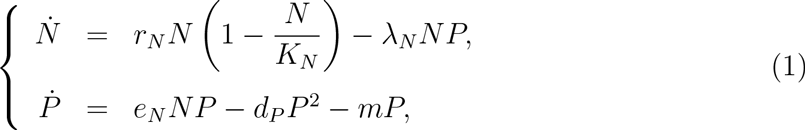

where the dot indicate derivatives with respect to time. The prey population grows logistically, with *r_N_* its intrinsic growth rate and *K_N_* its carrying capacity. We assume a linear functional response of the predator, with λ_*N*_ the capture rate of the predator. Predators grow proportionally to the amount of prey consumed *per* predator, with 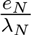 the conversion rate. They also negatively interfere, with a density dependent death rate *d_P_*, and die at a mortality rate *m*.

We also identified the Leslie-Gower model as another simple class of predator-prey model that exhibits negative density dependence. It assumes that prey have similar dynamics as in model (1) and that predators grow logistically with *r_P_* their intrinsic growth rate and a carrying capacity, *α_N_N*, proportional to the amount of prey consumed *per* predator. The model thus reads:

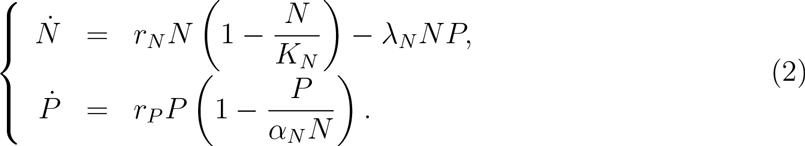

The predator equation differs from the one in model (1), which assumes that the reproductive rate of the predator population is proportional to its prey consumption rate, but its *per capita* predator reproductive rate is positively correlated with the number of prey consumed *per* predator, which has been identified as an important characteristic to be satisfied by predator-prey models (see *e.g*. [25,26]). Though model (1) has severe limitations when population becomes small [27], it has been used in many biological situations [28–31] and in various studies of population interactions [32,33].

### Introduction of an alternative prey with distraction effects

We extended models (1) and (2) to account for the presence of an alternative prey *A* in a situation where the predator establishes its predation activity on the two prey according to some trade-off (see [34] and references therein). Indeed, in nature, many prey species try to conceal or hide themselves to escape predation. To neutralize this strategy, predators can focus on specific cues emitted by a given prey species (visual, olfactory, auditory, etc.) to better locate and capture it. This is generically termed ‘formation of a search image’ by the predator [8,34]. Because any given organism can only process a limited amount of information in a given time period, forming a search image specific to a given prey type limits the efficiency in capturing other prey types: there is a time partitioning of predator’s attention for each of the prey species.

Search image models can provide significant impacts on predator-prey systems through the trade-off between capture rates of both prey represented here by parameter *q* ∈ (0,1) [35]. This parameter is termed the foraging ratio in this study. It represents the proportion of its time a predator focuses on prey *N*; correspondingly a proportion (1 − *q*) of predator’s time is dedicated to prey *A*. Introducing the foraging ratio in models (1) and (2) lead to two different models:

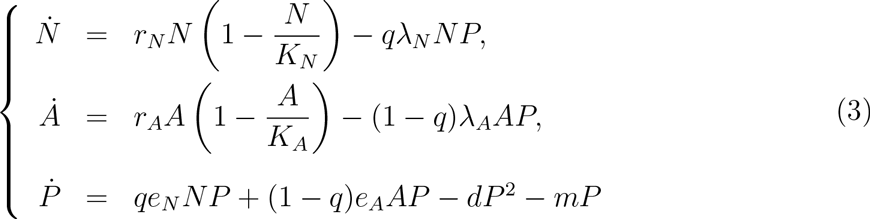

and

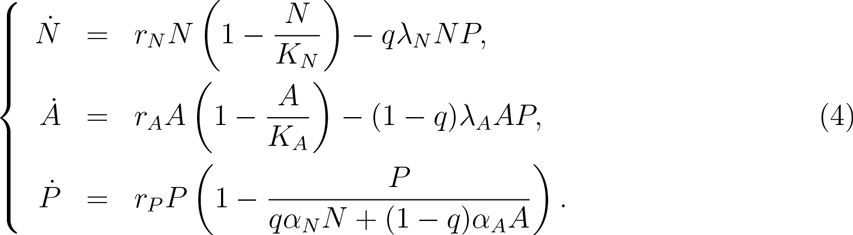

In both models, the dynamics of the alternative prey *A* are identical to prey *N*, with corresponding parameters *r_A_*, *K_A_* and λ_*A*_. An important difference with the single-prey models (1) and (2) is that, because of prey diversity and the trade-off resulting from the search image formation process, the capture rates of prey *N* and prey *A* are modulated by *q* and (1 − *q*) respectively. The reproduction term of predators *P* in model (3) and the carrying capacity of *P* in model (4) now depend on both prey. They are determined as the sum of the contributions of each prey: *qe_N_N* + (1 − *q*)*e_A_A* and *qα_N_N* + (1 − *q*)*α_A_A*, respectively. The foraging ratio *q* accounts for potentially time-varying predator preference for each prey: if *q* =1, *P* only forages for prey *N* whereas if *q* = 0, *P* concentrates on *A*. If *q* is strictly between 0 and 1, *P* has a mixed foraging behavior.

It should be noted that, even though it was motivated by cognitive processes, models (3) and (4) are also relevant in a spatially structured context, when prey species are segregated in space. If both prey inhabit habitats which do not overlap, but between which the predator can move freely, *q* and (1 − *q*) then represent the respective proportions of time spent by a predator in the *N* and *A* habitats (see *e.g*. investigations on coarse grained habitats in [17,36]). Such a behavior also refers to predator switching from one prey’s habitat to another depending on their respective density, which is known to stabilize predator-prey systems [37,38]. In this situation, the spatial segregation of prey imposes a partitioning of predator captures due to switching, similar to that brought about by prey search image formation.

Finally, let us comment on some of models (3) and (4) properties as compared to models (1) and (2), respectively. Suppose capture rates are equal and compare the one-prey models (1) and (2) with the respective two-prey models (3) and (4) with balanced foraging (*i.e. q* = 1/2). At the same overall number of prey in both scenarios (but spread over the two prey species for (3) and (4)), it is fairly easy to show that captures are more important in the one-prey environment (they are actually doubled because of the linear functional response). Conversely, adding the same number of an equivalent, but different, prey species halves the capture rate of the primary prey. This is a characteristic of the predator distraction effect: since prey are different, prey search image formation (or spatial segregation of prey) renders the predator unable to capture both species at a maximum rate at the same moment in time. All else equal, the distraction effect makes the predator less efficient in a multi-prey environment. Meisner *et al.* results on the distribution of the attack rates of *Aphidius ervi*, a natural enemy of aphids, provide a striking example of predator distraction (see Figure 2 in [12], albeit, strictly speaking, *A. ervi* is a parasitoid).

## Results

### Core of simple rules

Since the analytical expressions resulting from the analysis of model (4) are fairly simple, we chose to report the study of this model in the body of the results section, and detailed the analysis of model (3) in Appendix S3. Actually, the main property of the two models is that they are extensions of single-prey models may possess an equilibrium at which predators and prey coexist; when it exists, this equilibrium is globally stable. In what follows, we denote this single prey equilibrium 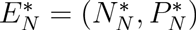.

For model (2), 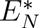 exists whatever the value of the carrying capacity; it reads:

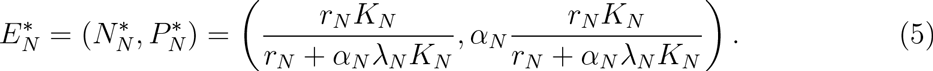

In the analysis of model (1), we assumed that *e_N_K_N_* > *m* which is the condition of existence of the coexistence equilibrium 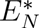.

Because one-predator-one-prey equilibrium values are important to predator-multi-prey systems, 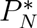, the realized predator equilibrium on prey *N*, will be given particular attention in what follows. We also introduce 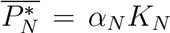 for model (2) (respectively 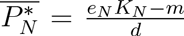 for model (1)), the ideal predator equilibrium on prey *N*, corresponding to the density a predator population would reach preying on a species *N* artificially maintained at its carrying capacity. Both the realized and ideal predator equilibria will be at the basis of simple rules aiming to characterize indirect effects.

### Fixed time partitioning strategy

The analysis of model (3) with fixed *q* shows that the dynamics always present a unique stable equilibrium, corresponding to the coexistence of the predator and one or both of its prey (Appendix S1.1). Indeed, a three-species coexistence equilibrium exists, provided that:

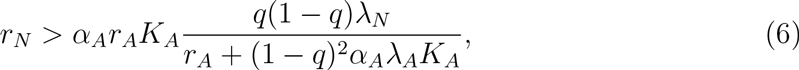

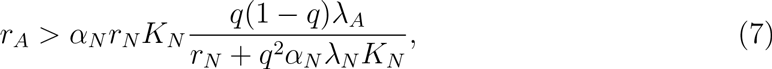

where (6) ensures the presence of prey *N* at equilibrium and (7) that of prey *A*; these conditions also ensure that the corresponding equilibrium is stable. Both conditions cannot be simultaneously transgressed (Appendix S1.1).

To elucidate the effects of predator foraging preferences, we studied the influence of *q* on the equilibrium of prey *N* and compared it with 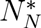, its value at equilibrium when *A* is absent. This leads to the following expression (Appendix S1.1):

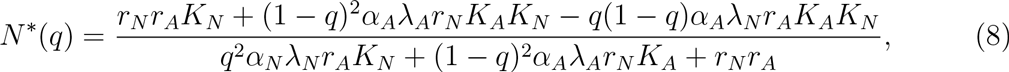

so that *N*\*(0) corresponds to the carrying capacity *K_N_* because *P* only forages for *A* and *N*\*(1) is equal to 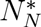 because *P* only forages for *N*. Two cases can then occur. Either the presence of the alternative prey is beneficial for *N* that reaches an equilibrium density comprised between 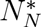 and *K_N_*, or detrimental to *N* that reaches an equilibrium density smaller than 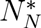. Actually, if 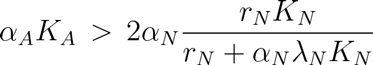 or, using the realized and ideal predator equilibrium definitions, if

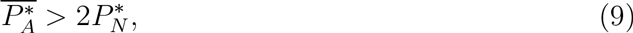

then prey *N* experiences negative effects for large values of *q*; otherwise, only positive effects can occur on prey *N* (Appendix S1.2). For large values of *α_A_* or *K_A_*, *i.e*. large 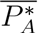, prey *N* can thus suffer negative effects. These can be strong enough to exclude *N*; that is when condition (6) is not satisfied. In a similar way, for low values of *α_N_,K_N_,r_N_* or large value of λ_*N*_, *i.e*. low 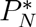, condition (9) tends to be more easily satisfied.

We get a symmetrical condition regarding the occurrence of detrimental effects from prey *N* on prey *A* for small values of *q*:

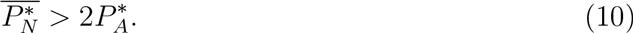

If this does not hold, only positive effects occur on prey *A*.

In the analysis of model (3), although forms (6), (7) and (8) differ, simple rules (9) and (10) are the same (Appendix S3). Thus the following discussion can be interpreted for both models (3) and (4).

Different effects on prey *N* were illustrated with prey which differ in parameters *e_i_* for model (3) (not shown) and in parameters *α_i_* for model (4) (Figure 1). In the first case, both prey contribute little to the predator population (low *e_I_* or *α_i_*) (Figure 1.A): *N*\*(*q*) stays above 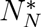 for all *q* ∈ [0,1]. Beneficial effects on both prey, *i.e*. apparent mutualism, are thus observed whatever the predator preference. The predator shares its foraging time between prey providing little to its population. As a consequence, the predator population does not take advantage of the low-quality prey and predator distraction effects dominate in the long term. In the second case, prey *A* quality has been increased (Figure 1.B) but *N* remains of moderate quality: *N*\*(*q*) remains above 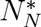 if the predator has a slight preference for *N*, but becomes smaller if the predator has a marked preference for *N*. Beneficial effects on prey *A* and detrimental effects on prey *N*, *i.e*. apparent predation, may be observed. This occurs because prey *A* is very beneficial to the predator because of a large *e_A_* or *α_A_* respectively, so that it increases the predator’s growth rate and thus predation pressure on prey *N* (the same would hold with a large *K_a_*). In the meantime, the high-quality prey benefits from predator distraction brought about by the low quality prey. In the third case, both prey are of high quality (Figure 1.C): both conditions (9) and (10) are satisfied so apparent predation and apparent mutualism can occur depending on *q* values. In such cases, apparent predation can however be strong enough to exclude one or the other prey.

In addition to these results, we showed that in both models at least one of the two prey always experiences beneficial effects (Appendix S1.3 and Appendix S3), preventing the occurrence of apparent competition and amensalism, so that at most one prey can suffer from negative indirect effects, resulting in apparent predation. Finally, temporal evolutions of the populations for model (4) have been computed to illustrate the effects of an introduction of prey *A*: because of predator distraction, *N* is indirectly favored in the short term, whereas it can benefit or suffer from the presence of *A* in the longer term (Figure S1, Appendix S2.4). Notice that in both the illustrated cases in Figure S1, the opposite effects occur as the multi-prey environment is profitable to the predator.

**Figure 1.**
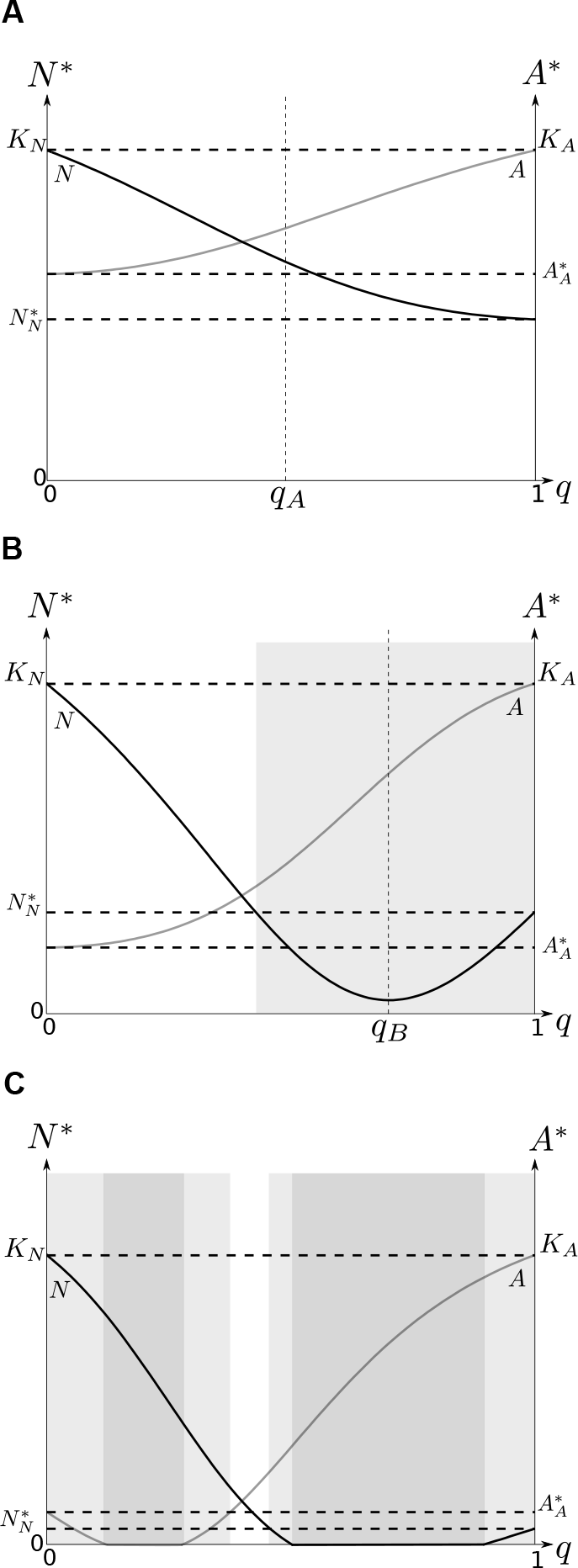
Dependence of N* and A on q. *N*\* and *A*\* are the densities of the primary prey and alternative prey at equilibrium, and are represented by black and gray curves, respectively. *q* is the preference of the predator. Most parameters are identical in all three subplots (*r_N_* = *r_A_* = 6; *K_N_* = *K_A_* = 3; λ_*N*_ = 3, λ_*A*_ = 2), except for parameters a which do vary (subplot A: *α_N_* = 0.7, *α_A_* = 0.6; subplot B: *α_N_* = 1.5, *α_A_* = 4; subplot C: *α_N_* = 12, *α_A_* = 8). The white areas correspond to values of *q* that induce apparent mutualism. The light gray areas correspond to values of *q* that induce apparent predation. The dark gray areas correspond to values of *q* that induce apparent predation with exclusion of one prey. *q_A_* and *q_B_* are particular values of the foraging ratio used in Appendix S1.4 to compute Figure S1.

### Adaptive time partitioning strategy

Adaptive foraging by predators has a major impact on food webs dynamics [40,41]. We thus considered the case in which the predator adapts its foraging ratio according to the actual prey densities. This resulted in predator switching between prey *N* and prey *A*: the predator chooses the most beneficial foraging ratio as a function of the environmental conditions [37,42]. We still used models (3) and (4) but contrary to the previous section, the foraging ratio *q* now varies in time: the predator population chooses *q* to maximize its growth rate. This actually leads to maximize *qe_N_N* + (1 − *q*)*e_A_A* and *qα_N_N* + (1 − *q*)*α_A_A* respectively, with respect to *q* at each moment in time. In fact, this instantaneous optimization is the limit of the dynamical fitness (*W*) maximization model

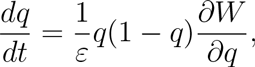

as behavioral adaptation (*q* adaptation) is fast compared to population dynamics, *i.e*. as ε tends to zero, and where fitness *W* is defined as the *per capita* growth rate of the predator (see *e.g*. [43]).

With an instantaneous *per capita* growth rate maximization behavior, the predator forages for the most profitable prey and switches to the other one when the former becomes scarce. In fact, we showed in Appendix S3.1 that in model (4) the predator forages for prey *N* and ignores prey *A* (*i.e. q* =1) if:

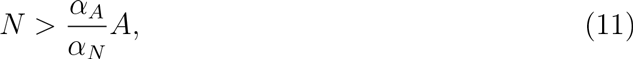

When inequality (11) is reversed, the predator switches to prey *A*. Thus a threshold separates the state space in two regions, where *q* = 0 or *q* = 1, respectively. In both regions, the non-predated prey follows logistic growth and the foraged predator-prey system follows model (2). If prey *N* has a density precisely equal to 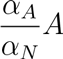, *i.e*. the system is on the threshold, the predator potentially forages for both prey. Yet, the relationship between *N* and *A* on this surface renders the predator dynamics independent from *q* and yields:

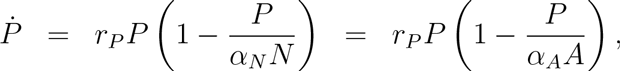

so that any *q* is optimal on the threshold.

Because the predator switches on the threshold, model (4) with adaptive *q* is discontinuous on this surface. It is actually the adaptive predator preferences in the remainder of the state space that impose specific dynamics and foraging ratio *q* onto this surface. This situation can be investigated using techniques developed for discontinuous differential equations [44,45].

Near the threshold, two situations can occur. On the one hand, trajectories on both sides of the threshold may be oriented in the same direction: solutions would then go through the threshold. On the other hand, trajectories may be oriented in opposite directions: the threshold is then either repulsive or attractive, depending on the orientation of the trajectories. In the former case, trajectories go away from the threshold and remain in one of the two regions, while in the latter case, trajectories converge to the threshold and stay on it, a phenomenon called sliding mode [44]. Two conditions ensure the existence of a sliding mode on the threshold and read (Appendix S3.2):

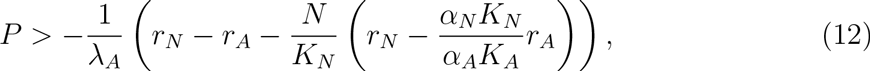

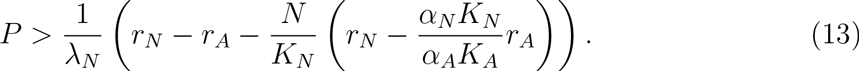

Since (12) and (13) cannot be simultaneously breached with *P* > 0, no part of the threshold is repulsive, so that the threshold is partitioned into two regions, one with sliding mode dynamics for high values of *P* when (12) and (13) are satisfied (region ***S***), and another where solutions go through. The analysis of the dynamics on the threshold can be performed using Filippov theory [44] but can also be presented in a more natural way. In region ***S***, both prey are of equal quality for the predator and a sliding mode will persist as long as this remains true. From (11) the threshold is invariant if:

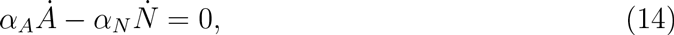

should hold true, so the foraging ratio in region ***S*** can be computed as:

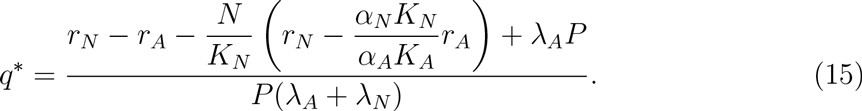

It can be shown that *q*\* necessarily belongs to [0,1] in region ***S***.

As *q*\* depends on state variables, the foraging ratio of the predator varies depending on the position of the system in region ***S***. Using the value of *q*\* given by (15), model (4) reduces to two equations in region ***S***:

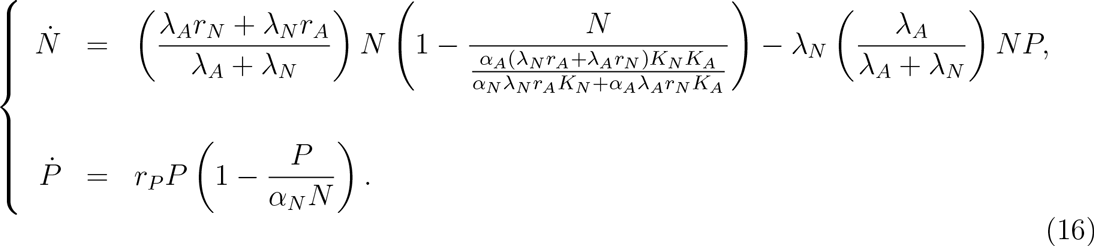

The complex behavior *q*\* of the predator actually renders the predator-prey system equivalent to a one-predator-one-prey Leslie-Gower model (2) which possesses a globally stable equilibrium:

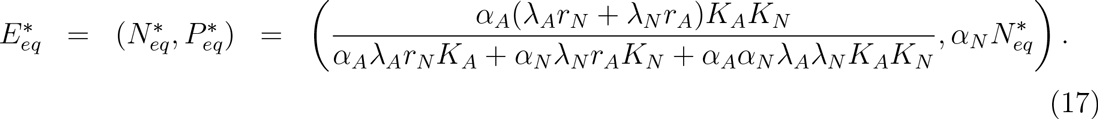

Yet, model (16) is only defined in region ***S*** of the threshold so that 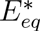 is an equilibrium of model (4) with adaptive *q* only if it lies in region ***S***, *i.e*. if it satisfies conditions (12) and (13). These existence conditions yield that *α_A_K_A_* > *α_N_N*\* and *α_N_K_N_* > *α_A_A*\*, *i.e*., using the realized and ideal predator equilibrium definitions,

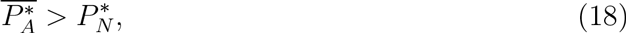

and

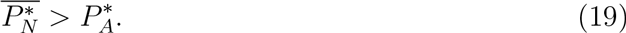

Straightforward computations actually show that both these existence conditions guarantee that 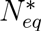 and 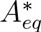 are larger than 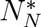 and 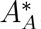, respectively, so that when it exists, 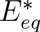 is characteristic of a positive indirect interaction between prey.

A similar analysis can also be done with model (3). On the threshold, the latter is reduced to a one-predator-one-prey Lotka-Volterra model (1), which admits a globally stable equilibrium on the threshold (Appendix S1). Actually, existence conditions of this latter are conditions (18) and (19) which also ensure positive indirect effects between prey. The following interpretation of these conditions thus holds for both models (3) and (4).

Figure 2 illustrates the two generic dynamics possible in model (4) with adaptive *q*: convergence to a long-term pure diet equilibrium (Figure 2.A) or to a mixed-diet equilibrium (Figure 2.B) with different ideal predator equilibrium values. These two situations can also be illustrated in model (3) depending on conditions (18) and (19). Actually, condition (18) means that prey *A* at carrying capacity is more valuable to the predator than prey *N* at the one-predator-one-prey equilibrium. Thus, if (18) holds, an adaptive predator cannot stick to a prey-*N*-only diet since at some point in time it would be advantageous to switch to prey *A*. If prey *A* is also more valuable at the single-prey equilibrium than prey *N* at the carrying capacity, *i.e*. if (19) is not verified, there is no advantage for the predator to switch back to prey *N*. In this situation, prey *N* benefits from the presence of *A* but has no effect on it: the predator ignores *N* and only forages for *A* (Figure 2.A). This one sided positive indirect effect is referred to as apparent commensalism of *N* with *A* [5]. Symmetrical results hold regarding prey *A*. However, if both (18) and (19) hold true, there are no pure diet equilibria so that only the mixed diet equilibrium 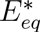 exists and is stable (Figure 2.B). In that case, long-term dynamics are of the apparent mutualism type.

One can additionally show that conditions (18) and (19) cannot simultaneously be breached (Appendix S3.3). This implies that, as in the fixed partitioning strategy, at least one prey always benefits from the time partitioning of the predator. However, in contrast with the results obtained for predators with a fixed preference for their prey, adaptive predator foraging only induces neutral or positive indirect effects between their prey, so apparent predation does not occur in this situation. Moreover, prey diversity is also profitable to the predator since one can easily show that 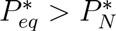 and 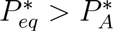 (Appendix S3.4). Temporal evolution of the populations in model (4) has also been computed to illustrate the effects on prey *N* of an introduction of prey *A*: positive effects induced by predator distraction in the short-term, and different outcomes in the long term (Figure S2, Appendix S1.5).

**Figure 2.**
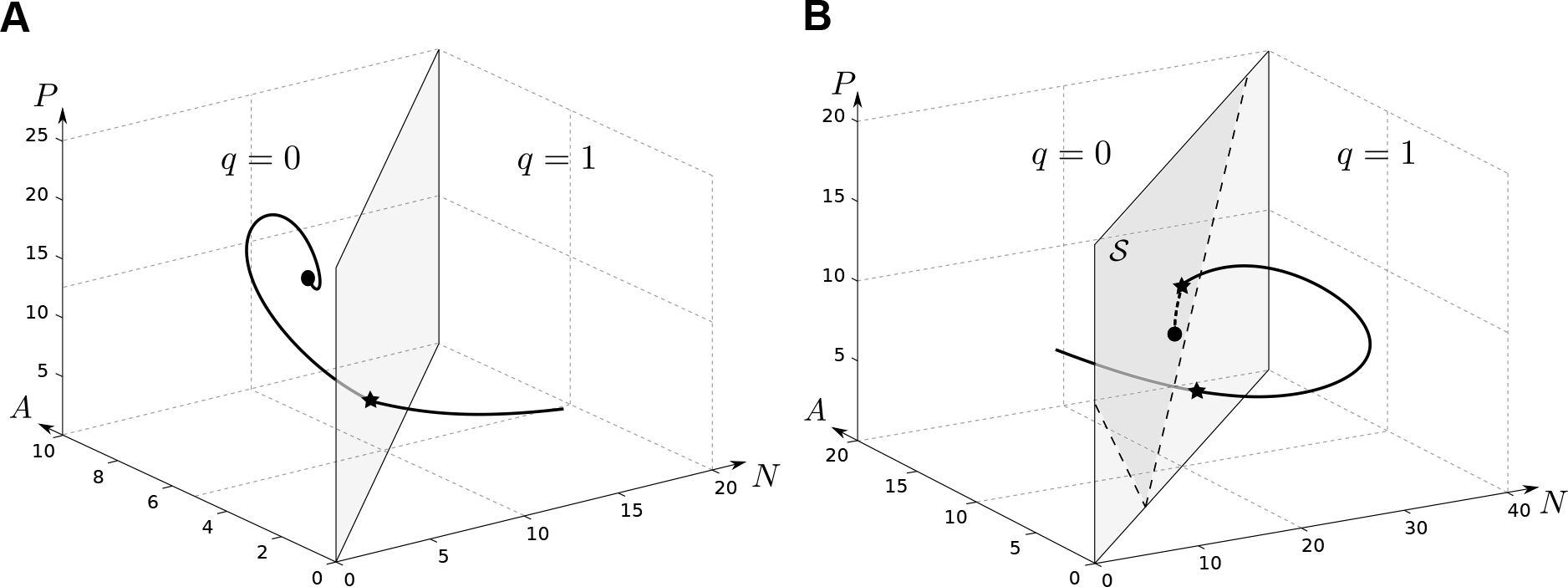
Evolution of the one-predator–two-prey system with adaptive preference. Most parameters are identical in all two subplots (*r_N_* = 12, *r_A_* = 15, *r_P_* = 12; λ_*N*_ = 1, λ_*A*_ = 1; *α_N_* = 0.5, *α_A_* = 1). In subplot *A*, prey *A* is the most profitable prey 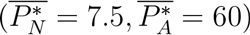 while in subplot B both prey are worthy to the predator 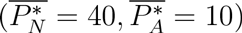. In both subplots, the light shaded surface figures the switching surface, while region S, delimited by the two dashed lines defined from sliding mode conditions (12) and (13), is represented in darker gray in subplot B only. Indeed, in subplot A condition (12) is only satisfied for large values of *P*, so that the trajectory is way below the sliding area and converges to the region where *q* = 0. In subplot B, the trajectory eventually converges to the sliding area, where *P* adopts a mixed diet, after having first crossed the threshold surface with condition (13) not satisfied.

### Foraging mode and positive indirect effects

To further explore the occurrence of indirect effects, we compared conditions when predators follow a fixed or an adaptive time partitioning strategy. We based our comparisons on the realized and ideal predator equilibrium values, in line with the previous work of Holt *et al.* [4]. We indeed showed that simple rules, as (9,10) or (18,19), can also characterize indirect effects in simple predator-prey systems influenced by complex mechanisms. Since these simple rules are observed for both Lotka-Volterra and Leslie-Gower models, they transcend model structures and can thus be considered as robust. We found that when the predator follows a fixed partitioning time strategy, apparent mutualism is guaranteed when both inequalities (9) and (10) are reversed, *i.e*. when 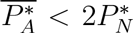 and 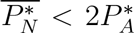. Otherwise, apparent mutualism or apparent predation, which can even lead to the exclusion of one or the other species, can happen depending on the value of *q* (Figure 1). When the predator follows an adaptive partitioning time strategy, apparent mutualism occurs when both conditions (18) and (19) are satisfied, *i.e*. when 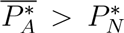 and 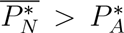. Otherwise, commensalism of one prey with the other happens. At this point one should notice the additional biological constraint that 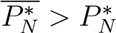 and 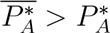.

The combination of these conditions yields three different generic situations determined in part by the realized predator equilibria on single prey 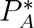 and 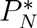: Both prey *A* and *N* may withstand the same, slightly or markedly different levels of predators at the predator-single-prey equilibrium. We illustrated these cases in Figure 3 by varying 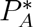 and keeping 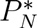 constant. In the first situation, *i.e*. 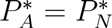, apparent mutualism occurs for all feasible 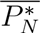 and 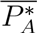 values when the predator adaptively forages (Figure 3.A, light gray region). In contrast, apparent mutualism is only guaranteed in the smaller portion of the parameter space located below 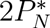 and 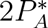 (Figure 3.A, dotted region) when the predator follows a fixed time partitioning strategy; otherwise, depending on the value of *q*, any indirect effect relevant to the fixed time partitioning strategy may occur. In the second situation, i.e. 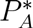 between 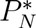 and 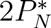, commensalism of *N* with *A* happens with the adaptive time partitioning strategy as 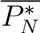 remains lower than 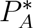 (Figure 3.B, dark gray region). A small region in which mutualism is guaranteed with the fixed time partitioning strategy, but does not occur with the adaptive one, also appears for low 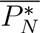 and 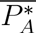 (Figure 3.B, dark gray and dotted area). In the third situation, as 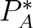 is larger than 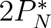, only the fixed time partitioning strategy qualitatively differs from the previous situation since no parameter combination guarantees the occurrence of mutualism anymore (Figure 3.C).

Although our conditions are slightly more complicated than the *P*\* rule, we were able to derive some simple rules ensuring the occurrence of unilateral and bilateral positive indirect effects driven by predator distraction, which can thus be generalized in the following situations. In a fixed time partitioning strategy, prey species should be much alike and correspond to small ideal predator equilibrium to experience apparent mutualism. Indeed, the realized predator equilibrium on one prey must be no larger than twice the realized equilibrium on the other, and the ideal predator equilibrium on one prey should stay below twice the realized predator equilibrium on the other. If one of the prey does not satisfy these rules, unilateral negative indirect effects can occur, leading to apparent predation. In an adaptive time partitioning strategy, the rule is even simpler since apparent mutualism happens when both ideal predator equilibria are larger than the largest realized predator equilibrium. However, if one of the prey does not satisfy these rules, apparent commensalism can occur. Thus, prey characterized by large 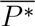 are more likely to experience apparent mutualism in an adaptive time partitioning strategy than in a fixed one. This outcome is reinforced by the fact that at most one prey can potentially suffer from negative indirect effects in the fixed time partitioning strategy, while prey are never penalized in an adaptive one. Such a discrepancy between foraging modes underscores the importance of characterizing predator foraging habits, especially in terms of adaptation to its prey environment.

**Figure 3.**
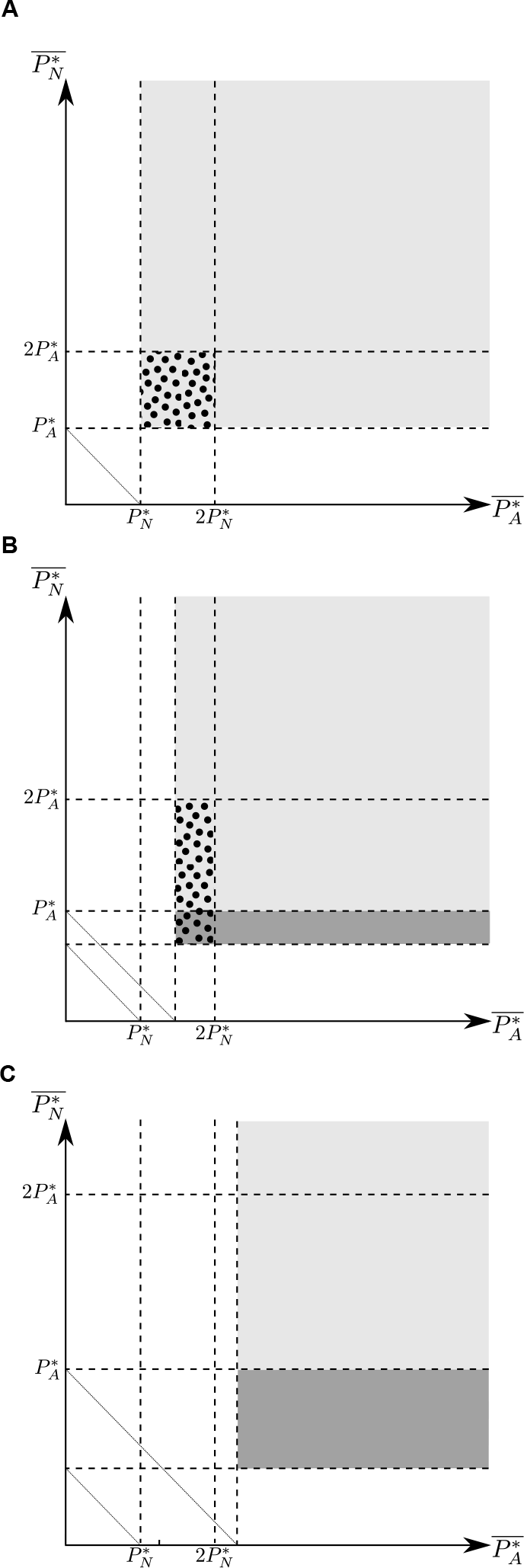
Occurrence of mutualism between prey depending on values of 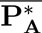 and 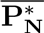. In subplot A, both prey can support the same level of predator at predator-single-prey equilibrium (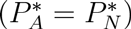); in subplot B, prey *A* is slightly more profitable than prey *N* at equilibrium (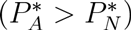) and in subplot C, prey *A* is markedly more profitable than prey *N* at equilibrium (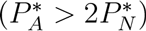). Four different regions are defined: unfeasible region (white area), ensured apparent mutualism with the fixed time partitioning strategy (dotted area), and apparent mutualism (light gray area) or apparent commensalism (dark gray area) with the adaptive time partitioning strategy.

## Discussion

The classical *P*\* rule disregards the fact that if a predator forages for many prey, it can have difficulties doing it at the same time. In some situations, such as when prey search image formation is needed or simply because of spatial segregation of prey species, the predator has to partition its time or its foraging effort, which releases predation pressure on each prey. This predator distraction effect directly induces positive indirect effects between prey species in the short term. However, predators’ numerical response may act concurrently on prey densities so that the combination of the two results in all likelihood in different types of indirect effects between prey species [17,18]. Most models developed to investigate indirect effects between prey species in predator-multi-prey systems conclude that apparent competition between prey is the rule in the long term, *e.g*. [4,7,36,46]. Nonetheless, it has been reported that apparent competition may be weakened in some specific situations, such as when the predator adaptively forages for its prey [47–50]. It has even been shown that mechanisms such as predator satiation, negative density dependence or adaptive foraging can reverse the indirect interactions and give rise to apparent mutualism between prey [19]. These situations make thus difficult the use of classical simple rules for describing complex predator-prey systems.

According to Holt’s original definition [7], “two species are in apparent competition whenever the presence of either species leads to a reduced population density for the other species at equilibrium”. In this study, we followed this definition, defining apparent mutualism and other indirect interactions, as apparent predation or apparent commensalism, as long term phenomena characterized by changes in equilibrium densities between the one-and two-prey scenarios. This approach has been the basis of the *P*\* rule which is built on the comparison of predator equilibria [4]. In the same way, we investigated indirect effects in the long term by comparing predator equilibria, whereas short term indirect effects linked to predator distraction have been evaluated as the instantaneous change caused by the presence of one prey on the other’s density. For that purpose, two classes of models based on different assumptions and structures have been explored. In both cases, similar indirect effects have been identified and characterized by identical rules based on realized and ideal predator equilibrium values. To go further in the exploration of indirect effects, we also compared simulations with either dynamical or instantaneous behavioral adaptation and found that both led to the same asymptotic results (simulations not shown). Drawing on these, we ensured the robustness of our conclusions and highlighted the relevance of simple rules for describing species interactions in community modules, even influenced by complex mechanisms.

In this paper, we indeed showed that positive indirect interactions can occur in the long term, once the predator population has numerically responded to the presence of different prey. In particular, we have obtained the intriguing result that higher predator densities do not necessarily lead to more prey suppression: predator distraction can counterbalance its numerical response and promote apparent mutualism between prey, and this is true whether predators forage adaptively or not. However, two main mechanisms can act against the occurrence of apparent mutualism. In the fixed time partitioning strategy, a prey with an ideal predator equilibrium larger than twice the realized predator equilibrium on the other prey may bring about a significant numerical response of the predator, and consequently affect negatively the other prey through apparent predation. In the adaptive time partitioning strategy, a prey with an ideal predator equilibrium smaller than the realized predator equilibrium on the other prey will eventually be completely ignored by the predator, leading to an apparent commensalism effect. Prey characteristics able to break apparent mutualism down are thus tighter in the adaptive scenario and also inhibit any negative indirect effect; this is because if adaptive foraging is beneficial to the predator, it also maximizes predator distraction and the consequent predator pressure release. Apparent mutualism is therefore more easily guaranteed in the adaptive rather than in the fixed case, while apparent predation only occurs in the latter case.

Our results complement previous findings on apparent mutualism reported in [7,18,19]. Although apparent mutualism is more frequent if predators are adaptive foragers, we showed that this is not compulsory: depending on prey characteristics, fixed time partitioning predators can indeed induce apparent mutualism for some, or all, values of the foraging ratio. Furthermore, predator distraction does not always weaken apparent competition equally on both prey. We thus identified situations when prey only experience unilateral positive effects so that apparent predation or apparent commensalism may occur. Actually, such indirect effects are known to be promoted by predatory behavior in the short term [17], and have recently been identified in the long term under the influence of predator saturation [51]. Our results showed that apparent predation can also occur with other mechanisms such as predator distraction and time partitioning strategies. They thus provide further underpinnings to the hypopredation phenomenon, a term which has been recently identified by [51] in the field of invasion biology: an invasive prey can actually be beneficial to a native prey by indirectly decreasing predation pressure, in contradiction with the hyperpredation concept, i.e. an increase in predation pressure [32]. Because adaptive behavior strongly weakens predators’ functional response on the focal prey, we did not found hyperpredation in the adaptive time partitioning strategy. In both strategies, we however emphasized on the fact that, at equilibrium, higher predator densities do not necessarily imply lower prey densities. As far as we know, this study is one of the first to precisely identify such phenomena in the long term. In fact, these phenomena contradict one of the main rationale underlying apparent competition theory, and may have important real-life consequences. Hence, depending on species interactions and predator behavior, different outcomes of multi-species predation can be expected. From the classical apparent competition evidenced in absence of time partitioning to apparent mutualism, its radical opposite shown in the present paper; we also produced intermediate outcomes in the form of apparent predation and apparent commensalism.

Positive interactions between species are increasingly recognized as important drivers of ecosystems structure and functioning, while a large part of the theory has so far concentrated on antagonistic interactions [52,53]. The picture is even a little more complicated since we have shown that complex interactions between individual behavior and population dynamics can turn an essentially antagonistic interaction into a unilateral or bilateral positive indirect one. These properties may be relevant in various applied areas, such as in the field of conservation biology: for instance, Halpern et al. [54] recommend a more thorough consideration of potential positive interactions in planning restoration or conservation programs in aquatic ecosystems. In a closer connexion with the present study, Sundararaj *et al.* [16] show that livestock populations can distract Asiatic lions from preying on native endangered chital deers, and advocate making the most of livestocks in conservation biology programs. As a matter of fact, lions’ numerical response could also be weakened by other factors than distraction, such as strong territorial behavior leading to direct interference, or large time scales required by the lions’ population to reach an equilibrium, leading to only observe short-term effects on prey. In fact, mechanisms promoting positive indirect effects are highly variable among systems and can result in a variety of indirect effects in nature. Identification of such mechanisms combined with extended simple rules are thus key to characterize indirect interaction in community modules, and could yield unifying principles in ecology based on equilibria comparison.

Our study can be specifically relevant to biological pest control program design. A large part of the experimental studies investigating indirect effects compare one-prey to two-(or more) prey scenarios (*e.g*. over 2/3 of the studies reviewed in [55] on predator-prey systems, see also [1]). This empirical approach is actually identical to the method we followed to analyze interactions in this paper. In this context, ideal and realized predator equilibria are simple indicators that could be experimentially estimated with single-prey and one-predator-single-prey systems, and then used to predict one-predator-multi-prey interactions. Since species behavior is known to alter biological pest control, mechanisms influencing species interactions, such as distraction effect, interference or spatial distribution, would need to be empirically identified prior to the start of experiments [56]. Experimental investigation into indirect effects could validate the combination of both indicators and identification of mechanisms for describing interactions in complex predator-prey systems. In turn, simple rules may help to ensure success of biological control programs or prevent potential failures by wisely choosing predators regarding their realized and ideal equilibria for example.

The present modelling study provided rationale supporting the empirical observation of Prasad and Snyder that “higher predator densities will not necessarily lead to improved pest control” [57]. A part of the effect reported in [57] is due to intraguild predation mechanisms, but distraction of generalist predators caused by the simultaneous presence of different prey is also acknowledged as an important driver of pest control disruption. This mechanism, originally put forward in [58], questions popular biological pest control practices. On the one hand, pest suppression cannot be determined by predator density alone; thus, against all odds, natural enemy density is not a reliable biological indicator of pest control. On the other hand, biological control programsbased on resource supplementation to biocontrol agents may turn counter-productive. For instance, providing alternative food to biological control agents can increase predator density, but also distract them from the targeted pest [10]. Similarly, shelters or banker plants [59] may also draw predators’ attention away from the targeted pest, which potentially disrupt pest control [57,60]. Because distraction effects have been reported in various biological control agents species (*e.g*. [10–13,57]), we believe that careful experimental studies should be conducted to assess the long term efficacy of resource supplementation in biological control programs.

## Acknowledgments

MTA was funded by a CJS grant from INRA. The authors acknowledge Nicolas Desneux for discussions on indirect interactions. Irma Mascio is thanked for her help in the English editing and Frank Hilker for his careful reading and comments on a previous version of this manuscript.

## Appendix

## S1 Analysis of model (4) with fixed time partitioning strategy

## S1.1 Equilibria and stability analysis

Model (4) with *q* fixed has the following boundary equilibria:

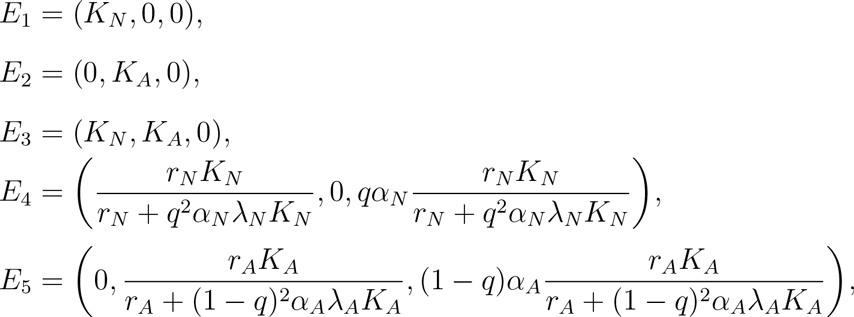

and a unique interior equilibrium: *E*\* = (*N*\*(*q*), *A*\*(*q*), *P*\*(*q*)) with

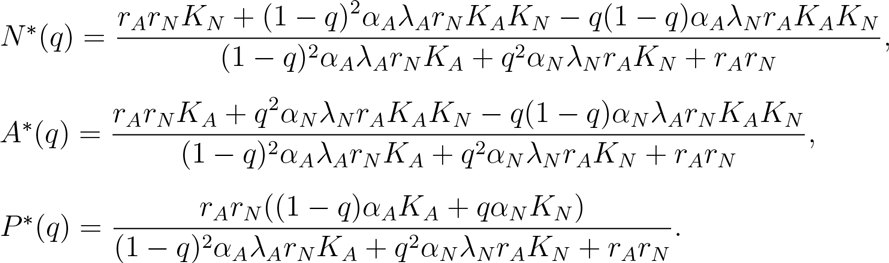

The equilibrium *E*\* exists if *N*\*(*q*), *A*\*(*q*), and *P*\*(*q*) are positive that is if both

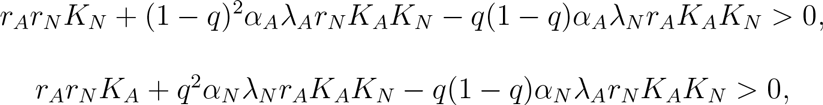

are satisfied. These conditions can be written as follows:

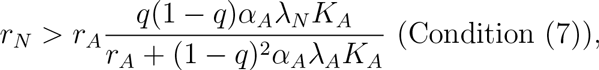

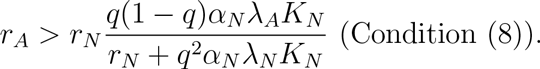

Both conditions cannot be transgressed simultaneously. Indeed, if they were, we could introduce *r_A_* obtained from the opposite of condition (8) into the opposite of condition (7) and obtain:

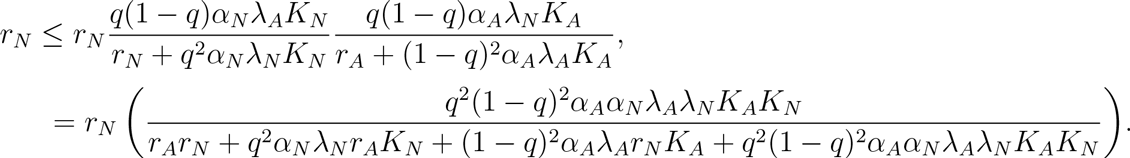

Since the numerator of the bracketed fraction is smaller than its denominator, this yields a contradiction and either condition (7) or (8) shall hold true.

The Jacobian matrix of model (4) at an equilibrium *Ē* = (*N̄*, *Ā*, *P̄*) is the following:

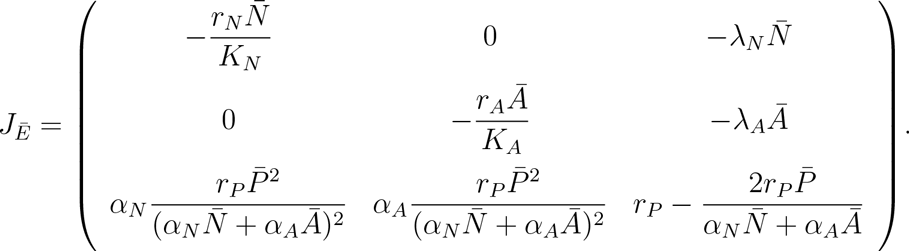

*E*_1_, *E*_2_ and *E*_3_ are unstable because *r_P_* is one of their eigenvalues. *E*_4_ is stable if condition (7) is not verified and E_5_ is stable if condition (8) is not fulfilled.

When both conditions (7) and (8) are satisfied, *E*\* exists and the characteristic polynomial of the Jacobian matrix is as follows:

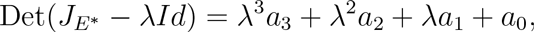

with 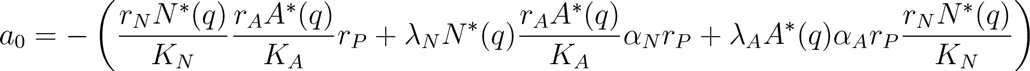, 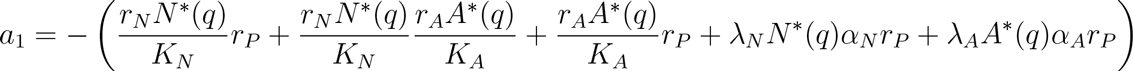, 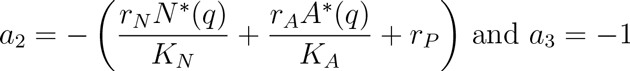

For this equilibrium to be stable, the Routh-Hurwitz criterion imposes that

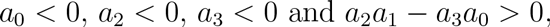

The three former conditions are verified when *E*\* exists, while we get for the latter:

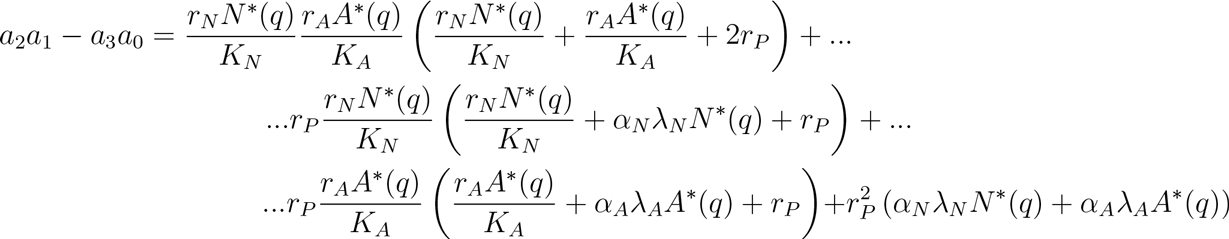

which is also positive if *E*\* exists. Then *E*\* is locally asymptotically stable.

## S1.2 Positive or negative effect on *N* of the introduction of *A*

We will study *N*\*(*q*) and show that it is always decreasing for *q* ∈ [0,1] or is decreasing to a minimum and then increasing to 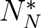. To do so, we need to consider *N*\*(*q*) as a function of *q* over ℝ, and not only restricted to the set [0,1] which is biologically sensible.

Several points need to be noticed for that:

- *N*\*(0) = *K_N_* > *N*\*(*q*) for all *q* ∈ (0, 1] (with 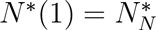).
- 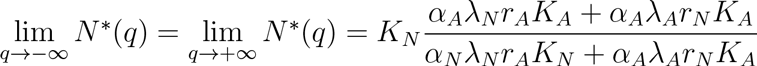.
- *N*\*(*q*) is continuous and has two extrema for *q* ∈ ℝ. Indeed

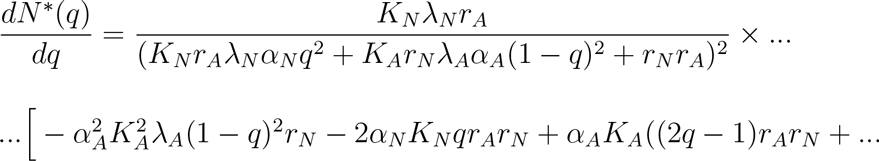

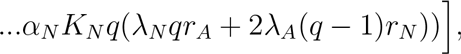

which cancels at the roots of the numerator, a 2^*nd*^ order polynomial. The two roots are real because 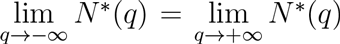 implies that, either *N*\*(*q*) is constant (which it is not), or it has at least an extremum.
- The derivative of *N*\* in *q* = 0 is negative:

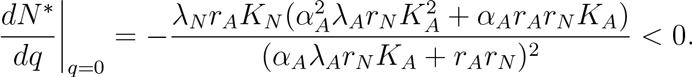
- The derivative of *N*\* in *q* =1 is as follows:

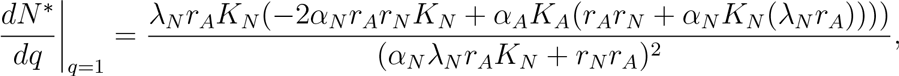

and is positive if 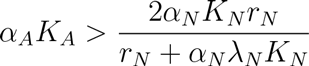 (condition (7)).

Since 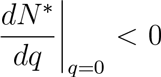, the first extremum *N*\*(*q*) has for *q* > 0, if it has one, must be a minimum. Therefore, three situations arise:

- *N*\*(*q*) has exactly one extremum for *q* ∈ [0,1), so that 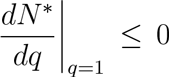. This implies that 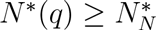 for all *q* ∈ [0, 1].
- *N*\*(*q*) has exactly one extremum for *q* ∈ (0, 1), which needs to be a minimum, having 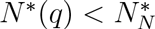, and no maximum in *q* = 1. This is equivalent to having 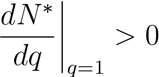.
- *N*\*(*q*) has two extrema for *q* ∈ (0, 1], a minimum then a maximum, so that 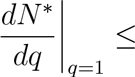 0. Since *N*\*(*q*) has no more than two extrema over ℝ, this implies that *N*\*(*q*) is always decreasing for *q* outside the interval so that 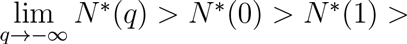 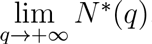 which leads to a contradiction.

## S1.3 At least one prey always benefits from the time partitioning

If *q* = 0 or *q* = 1, one of the prey is ignored so that it benefits from a positive indirect effect from the other prey.

Consider *P*\*(*q*) = *qα_N_N*\*(*q*) + (1 − *q*)*α_A_A*\*(*q*) the equilibrium of the predator *P* for *q* ∈ (0,1) from the *Ṗ* = 0 equation. Suppose that *P*\*(1) = *α_N_N*\*(1) ≥ *P*\*(0) = *α_A_A*\*(0). Suppose moreover that there exists *q* such that *N*\*(1) ≥ *N*\*(*q*) > 0 and *A*\*(0) ≥ *A*\*(*q*), which means that both prey experience detrimental effects for this *q* value. As consequences,

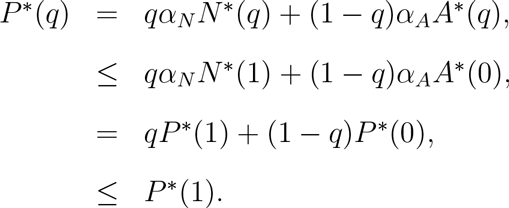

Since *N*\*(*q*) = 0, from the *Ṅ* ≠ 0 equation, 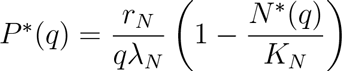. Since 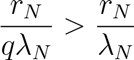 and 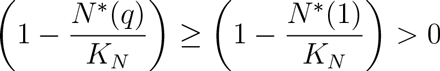, we have:

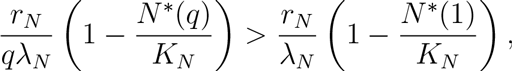
which means that *P*\*(*q*) > *P*\*(1). That contradicts the initial assumptions.

Because both prey play a symmetric role in the equations, similar contradictions occur if we assume *P*\*(0) ≥ *P*\*(1) and follow a symmetrical reasoning. Thus indirect effects without extinction are always positive for at least one of the prey.

The only roadblock to the preceding proof occurs when either *N*\*(*q*) = 0 or *A*\*(*q*) = 0.

Suppose *N*\*(*q*) = 0 and *A*\*(*q*) ≤ *A*\*(0). We have:

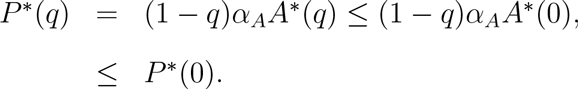

Then, since *A*\*(*q*) > 0 because both species cannot go extinct simultaneously, we have, from the 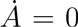 equation, 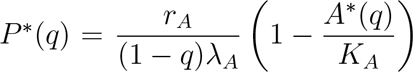. Since 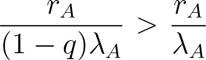 and 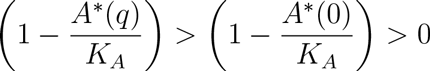, we have:

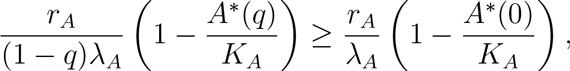

which means that *P*\*(*q*) > *P*\*(0). That contradicts the initial assumptions.

A symmetrical reasoning can be held in the case where *A*\*(*q*) = 0, which concludes the proof that at least one prey always benefits from the predator’s time partitioning.

## S1.4 Temporal evolution of the populations

Temporal dynamics of prey *N* were computed (Figure S1) to illustrate the effects on *N* of the introduction of *A*. Before the introduction, the predator only forages for *N* (*q* = 1) then partitions its foraging time as *A* is introduced, which relaxes its pressure on the primary prey in the short term. Beneficial effects in the short term are thus due to a distraction effect on the predator. Depending on how the alternative prey improves the growth rate of the predator according to the *q* values, long-term positive indirect effects can occur on prey *N* if both prey have low *α_i_* (subplot A) or long-term negative indirect effects if the alternative prey is such that *α_A_* ⪢ *α_N_* (subplot B).

**Figure S1.**
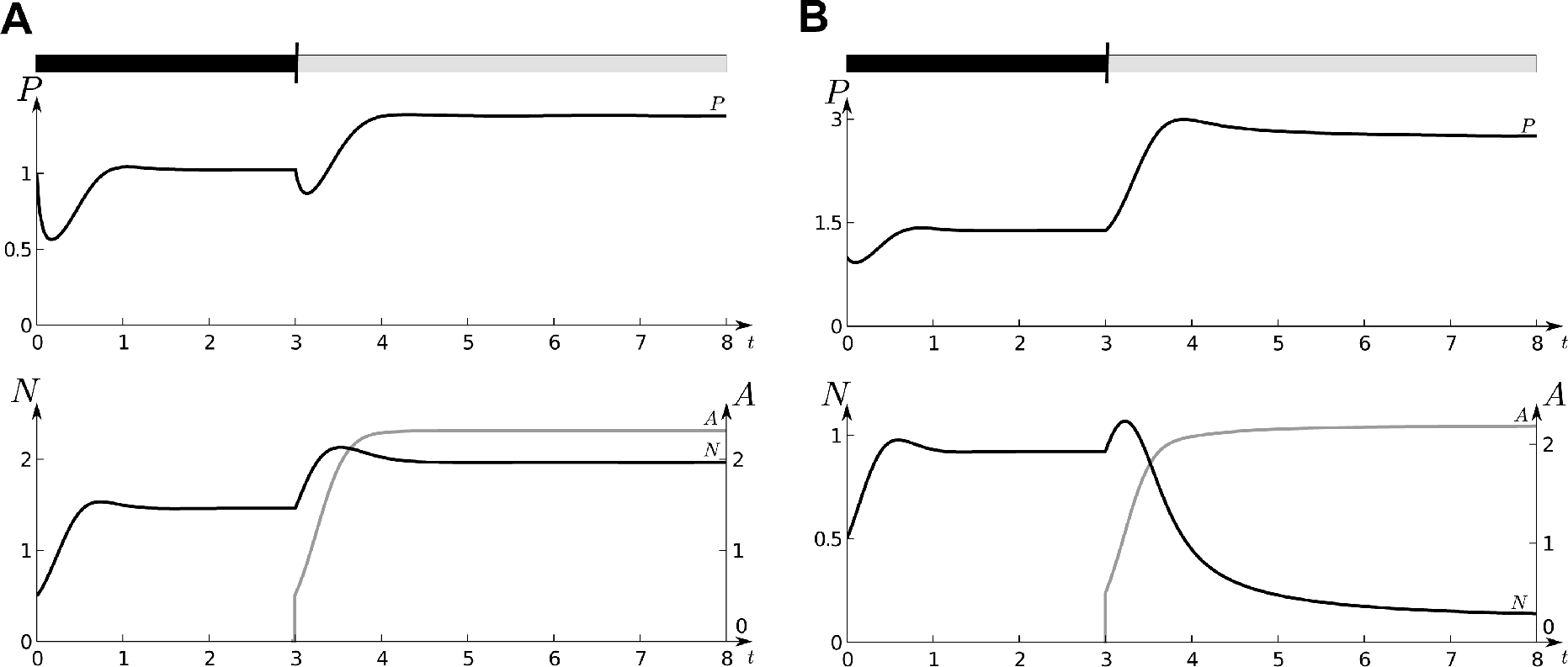
Effects of the introduction of an alternative prey on a one-predator-one-prey system with fixed time partitioning strategy. All parameters are the ones defined for Figure 1.A and Figure 1.B, respectively. *q* = *q_A_* is defined in Figure 1.A, and *q* = *q_B_* in Figure 1.B. Initially, *q* = 1, during a timespan represented by the black rectangular shape, *A* is then introduced (black trait above the figures) and the predator adopts a fixed *q* (timespan represented by the grey rectangular shape).

## S2 Analysis of model (4) with adaptive time partitioning strategy

## S2.1 Partition of the state-space

The predator optimally forages for its prey by maximizing its intrinsic growth rate, which is equivalent to:

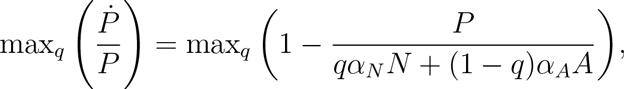

which is then achieved for the *q* value that maximizes *K*(*q*) = *qα_N_N* + (1 − *q*)*α_A_A*. We have:

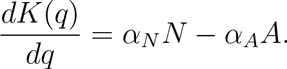

So the predator maximizes its intrinsic growth rate by:

- only foraging for *A* if 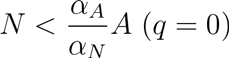 because *K*(*q*) is a decreasing function of *q*.
- only foraging for *N* if 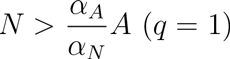 because *K*(*q*) is an increasing function of *q*.
- adopting some mixed diet if 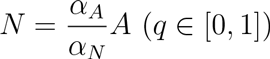 because *K*(*q*) is constant.

## S2.2 Sliding mode conditions

Let 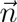 be the normal vector to the surface 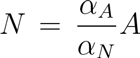, oriented from the region where *q* = 0 toward the region where *q* =1.

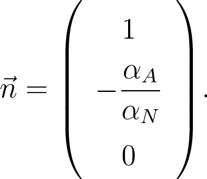

Let denote 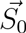 and 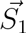 the right-hand sides of model (4) when *q* = 0 and *q* =1, respectively. Sliding mode is observed on the surface 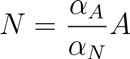 if:

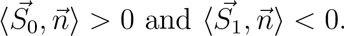

On the one hand:

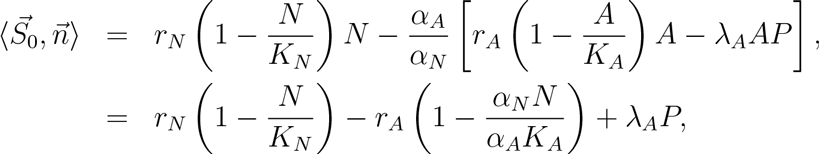

where the equality 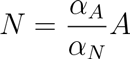 has been used. Then the condition 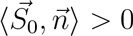 is equivalent to:

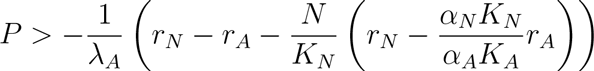

On the other hand:

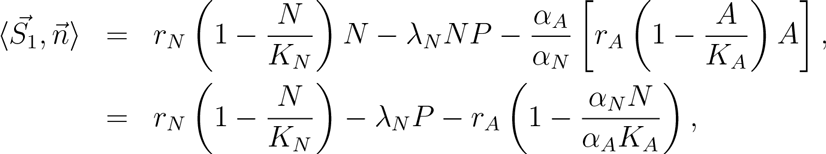

where the equality 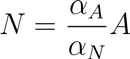 has been used. Then the condition 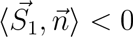 is equivalent to:

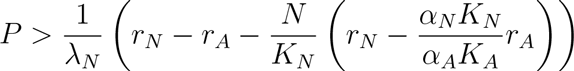

## S2.3 At least one prey benefits from the time partitioning

Here we show that either condition (19), (20) or both hold true, so that at least one of the two prey benefits from the adaptive time partitioning.

Suppose that both (19) and (20) are breached so that:

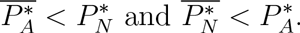

Recall that we necessarily have the feasibility conditions 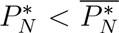 and 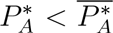. Thus, using both breached conditions and the first feasibility condition, we have:

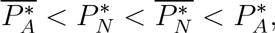

which contradicts the second feasibility condition.

## S2.4 The predator always benefits from its mixed diet

The predator benefits from its mixed diet if 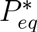 is larger than 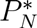 and 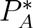, which leads to:

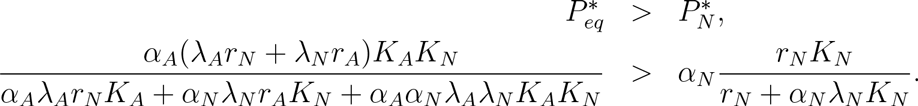

We develop and simplify the two members of the inequality:

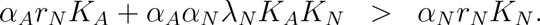

We then factorize the inequality and obtain:

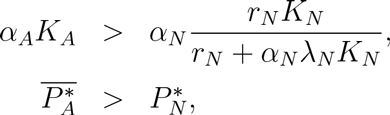

which results on condition (19).

We get the symmetrical condition (20) regarding

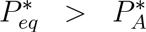

which reads:

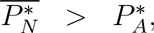

Thus, if 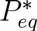 exists, *i.e*. if the predator adopts a mixed diet, it always reaches a higher density at equilibrium than in pure diets.

## S2.5 Temporal evolution of the populations

We illustrated the cases where condition (19) holds true but (20) is not satisfied (Figure S2.A), and where both conditions are satisfied (Figure S2.B). In Figure S2.A, a one-predator-low-quality-prey system is considered; only prey *N* is present at the beginning of the simulation; thereafter a better, but still low-quality, prey *A* is introduced. In this type of situation, condition (19) holds true, but (20) is not satisfied, hence 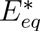 is not an equilibrium of the model. At first, the system is not influenced by *A* because it is of low density. The predator ignores *A* until the system reaches the threshold, because of the growth of *A*. As this occurs, the predator switches from *N* to *A*, which has just become the most profitable prey. Since *N* is a lower-quality prey than *A*, it is ignored by the predator in the long-term and reaches its carrying capacity, whereas *P* focuses on *A* (*q* = 0). *N* experiences short-term neutral effects followed by positive effects due to the switching of the predator to the higher-quality prey. In the long term, the presence of prey *A* thus releases predation from prey *N* which tends to its carrying capacity. Long-term dynamics of prey *A* appear unaffected by the presence of prey *N* since the predator behaves as if the latter was missing (apparent commensalism). A symmetric situation takes place for *A* and *N* if (20) holds true, but (19) is not satisfied. Thus both prey can reversely experience commensalism, depending on which is the most profitable for the predator. Figure S2.B illustrates a similar scenario in which both prey are of high-quality so that (19) and (20) hold and 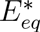 is the only stable equilibrium of the system. At first, prey *N* experiences short-term positive indirect effects thanks to a distraction effect: the predator immediately switches from *N* to *A* which is of high density. However, because *N* is also valuable, the system eventually reaches region ***S*** and ultimately converges towards the co-existence equilibrium 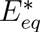. Therefore, both prey experience long-term positive effects by reaching equilibrium values that are higher than the ones corresponding to one-prey only situations (apparent mutualism).

## S3 Analysis of model (1) with an additional prey

## S3.2 Analysis of model (1)

In what follows, we will consider that predators can persist in the system with only one prey, *i.e. e_N_K_N_* > *m*. Model (1) thus admits a unique globally stable equilibrium at which predators and prey coexist:

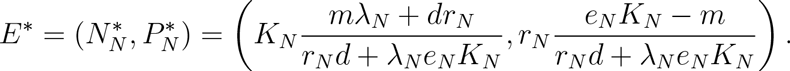

We consider that 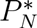 is the realized predator equilibrium on prey *N*, and we introduce 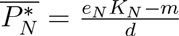, the ideal predator equilibrium on prey *N*. We do not detail the methodology that leads to the main results in that follows, since the assumptions and mathematical steps remain the same that the ones we have used to identify indirect effects with the Leslie-Gower model.

**Figure S2.**
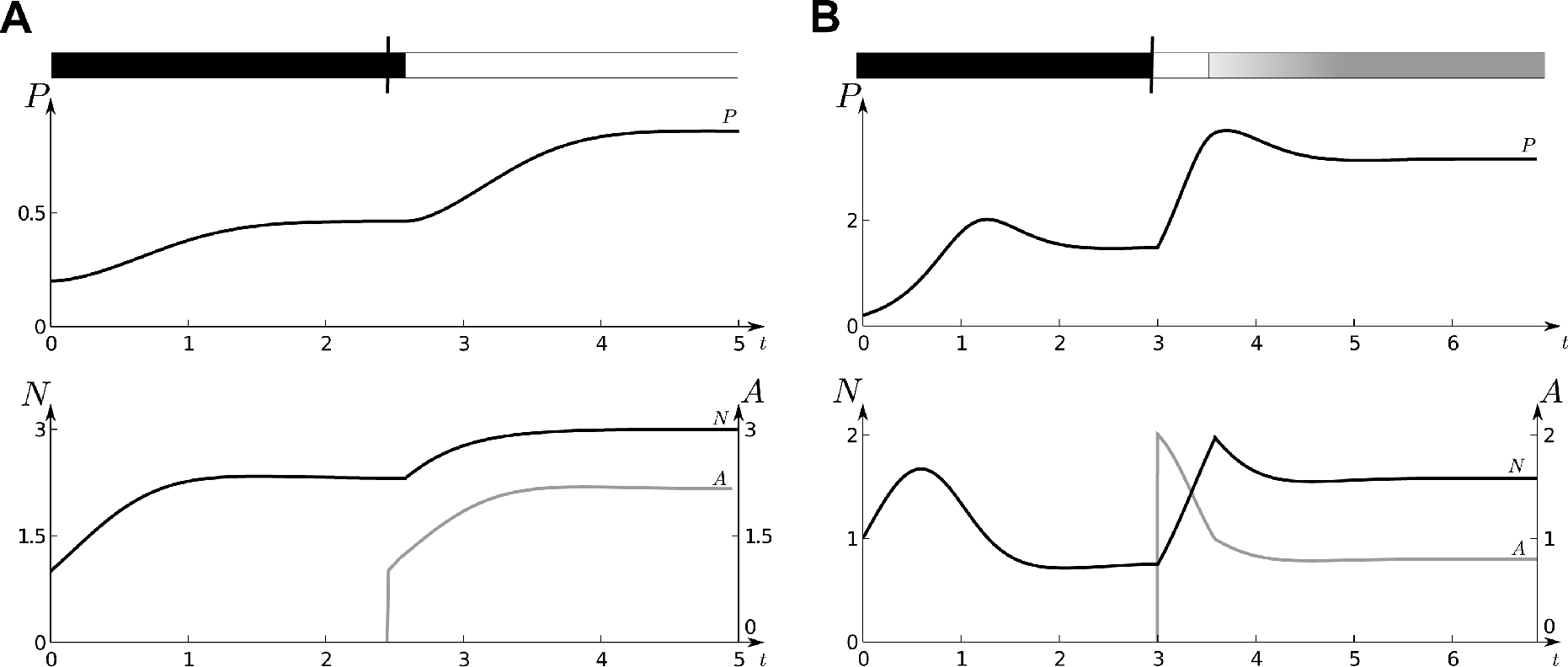
Effects of the introduction of an alternative prey on a one-predator-one-prey system with adaptive time partitioning strategy (r_N_ = r_A_ = 6; K_N_ = K_A_ = 3; λ_N_ = 3, λ_A_ = 2). Initially, *q* =1, during a timespan represented by the black rectangular shape, A is then introduced (black trait above the figures). In subplot A (*α_N_* = 0.2, *α_A_* = 0.4), *P* switches from *N* to *A* when (9) is not satisfied anymore (*q* = 0 represented by the white rectangular shape). In subplot B (*α_N_* = 2, *α_A_* = 4), *P* instantaneously switches from *N* to *A* (*q* = 0 represented by the white rectangular shape) then reaches the switching surface and stays on it (*q* ∈ [0,1], gradient of the gray rectangular shape).

## S3.2 A one-predator–two-prey model

To ease the reading, we recall model the Lotka Volterra one predator two-prey model (3):

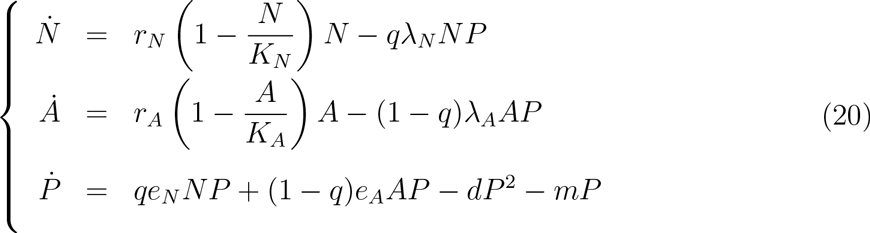

## S3.3 Fixed time partitioning strategy

If *qe_N_K_N_* + (1 − *q*)*e_A_K_A_* − *m* > 0, model (3) admits a unique stable equilibrium *E*\* = (*N*\*(*q*), *A*\*(*q*), *P*\*(*q*)) with:

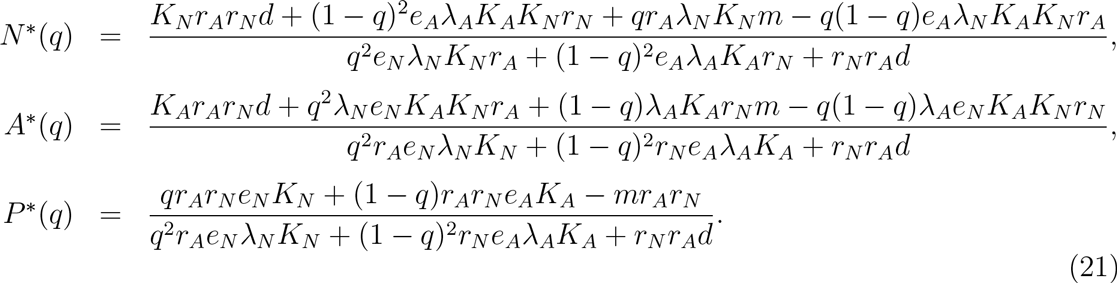

We studied the influence of *q* on the equilibrium of prey *N* and compared it with 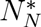. Following the methodology introduced in Appendix S1.2, we concluded that *A* has a positive effect on *N* for any value of *q* if:

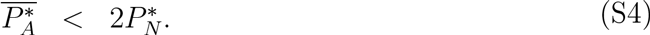

We get a symmetrical condition regarding the occurrence of positive effects from prey *N* on prey *A*, which leads to the occurrence of apparent mutualism if the following conditions are satisfied:

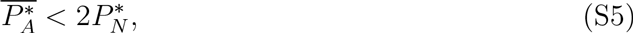

and

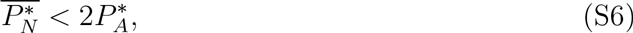

As demonstrated in Appendix S1.3, at least one prey always benefits from the time partitioning: the indirect effects range from apparent predation and punctual apparent commensalism to apparent mutualism.

## S3.4 Adaptive time partitioning strategy

The predator can choose *q* to maximize its growth rate. The predator thus forages for prey *N* and ignores *A* (*i.e. q* = 1) if:

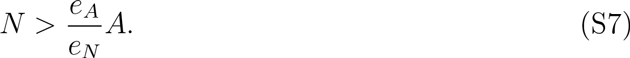

If the inequality (S7) is reversed, the predator switches to prey *A* (*q* = 0). If prey *N* has a density equal to 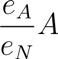, the system is at a threshold separating the two regions where *q* = 0 and *q* = 1, respectively. On the threshold, the system admits a unique stable equilibrium 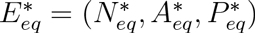 with

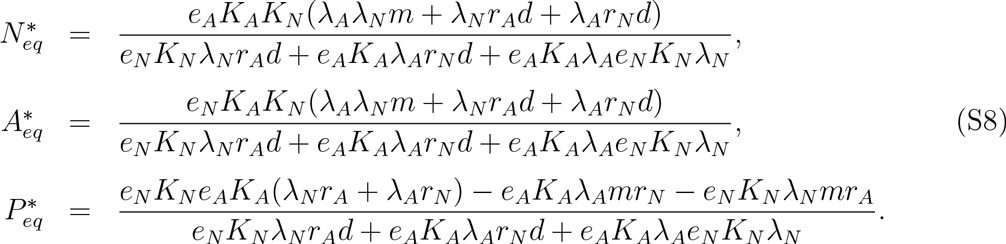

The existence conditions of 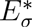 yield that:

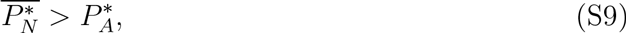

and

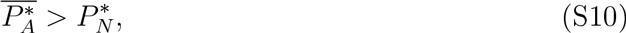

These conditions guarantee that 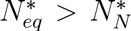 and 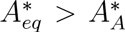, so both prey experience apparent mutualism when the predator adopts a mixed diet (*q* ∈ [0,1]). If condition (S9) is not satisfied, prey *N* benefits from the presence of prey *A* but has no effect on it, which refers to apparent commensalism. The symmetrical results hold regarding prey *A*.

